# Development and Function of Ovarian Lymphatic Vasculature

**DOI:** 10.64898/2026.05.26.727976

**Authors:** Subhasri Biswas, Lijuan Chen, Julia Skalka, Lijun Xia, R. Sathish Srinivasan, Michael B. Stout, Xin Geng

## Abstract

Lymphatic vessels play essential roles in interstitial fluid clearance and immune cell trafficking. Recent studies have uncovered previously unrecognized, organ-specific functions of the lymphatic vasculature in the meninges, heart, kidney, and lungs. However, the development and physiological significance of the ovarian lymphatic network remain poorly defined. To address this gap, we performed high-resolution three-dimensional imaging of ovarian lymphatic vasculature in mice. Our results reveal that the ovarian lymphatic network comprises lymphatic capillaries and valve-containing pre-collectors and the ovarian lymphatic vasculature arises from pre-existing lymphatic vessels in the broad ligament starting at postnatal day 5. Furthermore, we demonstrate that the VEGF-C/VEGFR-3 signaling is essential for ovarian lymphatic development and *Vegfr*3 insufficiency impairs ovarian lymphangiogenesis, leading to fibrosis and premature ovarian aging. Together, these findings identify the lymphatic vasculature as a critical component of the ovarian microenvironment and establish a causal link between lymphatic insufficiency and accelerated ovarian aging.

## INTRODUCTION

The lymphatic vasculature plays essential roles in fluid homeostasis, immune surveillance, and clearance of cellular debris, with growing evidence supporting organ-specific functional specializations beyond these canonical roles (1, 2). Meningeal lymphatics facilitate cerebrospinal fluid (CSF) drainage and metabolic waste clearance from the central nervous system (CNS), and their age-related decline accelerates neurodegeneration and cognitive impairment (3–5). Renal lymphatics regulate interstitial fluid homeostasis and inflammatory responses during kidney injury (6) . Pulmonary lymphatics are essential for edema clearance and immune regulation in the lung, and their dysfunction contributes to the pathogenesis of pulmonary hypertension and post-transplant complications (7, 8). Cardiac lymphatics maintain interstitial fluid homeostasis and immune cell trafficking within the myocardium and support cardiomyocyte proliferation and survival during both heart development and ischemic cardiac disease (9, 10). These organ-specific functions have driven the development of targeted therapeutic strategies to enhance lymphatic drainage across disease contexts. In the CNS, intracisternal delivery of VEGF-C enhances meningeal lymphatic drainage and improve cognitive outcomes in preclinical models of neurodegeneration, while activation of the mechanosensitive ion channel Piezo1 via the agonist Yoda1 restores CSF outflow in models of craniosynostosis and aging (3, 11–13). In the heart, stimulation of cardiac lymphangiogenesis through VEGF-C/VEGFR-3 signaling reduces myocardial edema, resolves inflammation and improves cardiac function following myocardial infarction (14, 15). Collectively, these findings underscore that lymphatic vessels are active regulators of organ homeostasis and disease, highlighting their therapeutic potential and underscoring the need to define lymphatic structure and function in organs where their roles remain poorly understood.

The ovary is a highly dynamic organ that undergoes continuous remodeling throughout reproductive life. During each cycle, follicle-stimulating hormone recruits a cohort of follicles, yet typically only one reaches ovulation while the remainder undergo atresia. Ovulation is a tightly regulated process involving immune cell recruitment and extracellular matrix remodeling to enable follicular rupture and oocyte release, followed by rapid formation of the highly vascularized corpus luteum (16). In the absence of fertilization, the corpus luteum regresses and is cleared by phagocytic cells (17). These recurring cycles generate a substantial burden of cellular debris and extracellular matrix turnover, necessitating efficient clearance to maintain ovarian architecture and function. Impaired clearance of interstitial fluid, macromolecules, and inflammatory mediators promotes stromal fibrosis, while increased fibrotic remodeling and tissue stiffness disrupt folliculogenesis and contribute to ovarian insufficiency (16, 18, 19). Notably, the ovary is among the earliest organs to exhibit functional decline, with fertility decreasing after age 35 followed by menopause at 51 years of age on average (20). Beyond reproduction, ovarian endocrine function is essential for maintaining bone, cardiovascular, and neurological health, underscoring the systemic consequences of ovarian functional declines (21, 22). In some women, ovarian aging is accelerated by identifiable insults including chemotherapy, radiation, autoimmune disease and genetic mutation, or occurs through incompletely understood mechanisms, as in idiopathic premature ovarian insufficiency (POI) (23–26).

Given the established roles of lymphatic vessels in waste clearance and immune regulation, and the ovary’s substantial demands in both processes, a functional ovarian lymphatic network is likely essential for tissue homeostasis. Disruption of lymphatic function has been shown to perpetuate chronic inflammation, impair tissue repair and alter immune surveillance across multiple organ systems, raising the possibility that similar mechanisms contribute to follicular atresia, stromal fibrosis and the dysregulated immune milieu observed in ovarian aging and premature ovarian decline (27–30). Although several studies have examined ovarian lymphatics (31–34), their developmental origin, three-dimensional organization, and molecular regulation remain incompletely defined, and the contribution of lymphatic dysfunction to ovarian aging is unknown.

In the present study, we address these questions through a systematic analysis of ovarian lymphatic development and function in mice. The mouse ovary is a well-established model for human ovarian disease due to conserved anatomical and physiological features, including folliculogenesis and hormonal regulation via the hypothalamic-pituitary-gonadal (HPG) axis (35–37). Mouse models have provided critical insights into conditions such as POI, polycystic ovary syndrome (PCOS), granulosa cell tumors and age-related ovarian reserve decline (35). Although differences exist, including a shorter estrous cycle, polyovulation and the absence of menstruation, the mouse ovary remains the gold standard preclinical model. Using high-resolution three-dimensional imaging, genetic lineage tracing, scanning electron microscopy, RNAscope and a model of lymphatic insufficiency, we define the stepwise postnatal development of ovarian lymphatics, establish that these vessels arise from pre-existing lymphatic endothelium, identify VEGF-C/VEGFR-3 as the critical signaling pathway for ovarian lymphangiogenesis, and demonstrate that chronic lymphatic insufficiency leads to premature ovarian aging accompanied by immune infiltration and fibrosis. Together, these findings establish the lymphatic vasculature as a previously underappreciated but essential component of the ovarian microenvironment and implicate lymphatic dysfunction as a potential driver of ovarian functional declines.

## RESULTS

### The ovarian lymphatic vasculature develops postnatally in a stepwise manner

To define the spatial and temporal development of the ovarian lymphatic vasculature, we performed high-resolution three-dimensional imaging of whole-mount ovaries from postnatal day 4 (P4) to P10 and P15 (Figure 1A-C and data not shown). Immunofluorescence staining for lymphatic markers PROX1, LYVE1, and VEGFR-3, together with pan-endothelial marker CD31, revealed an absence of lymphatic vessels in P4 ovaries (data not shown). In contrast, lymphatic vessels were detected in a subset of P5 ovaries, indicating lymphatic vessels first enter the ovary through the hilum at P5, at a stage when a dense blood vascular network is already established (**Figure 1A**). Upon entry, LYVE1⁺ lymphatic vessels closely associate with CD31⁺ ovarian veins and arteries, as well as TUJ1⁺ peripheral nerves, forming organized neurovascular-lymphatic bundles that extended into both the ovary and adjacent uterus (**Figure 1D**). Following hilum entry, lymphatic vessels progressively extend along the anterior–posterior axis of the ovary. By P10, lymphatic vessels reach the mid-ovarian region, and by P15 they form an organized network in close proximity to developing follicles (**Figure 1B**). As development progresses, additional lymphatic sprouts emerged not only from the hilum entry site but also from pre-existing intraovarian vessels, indicating that secondary sprouting contributes to the expansion and elaboration of the lymphatic network (**Figure 1B**). Low levels of PROX1 expression were also detected in oocytes as previously reported (33) (data not shown). The sequential steps of ovarian lymphatic vascular development are summarized in Figure 1C.

**Figure 1.**
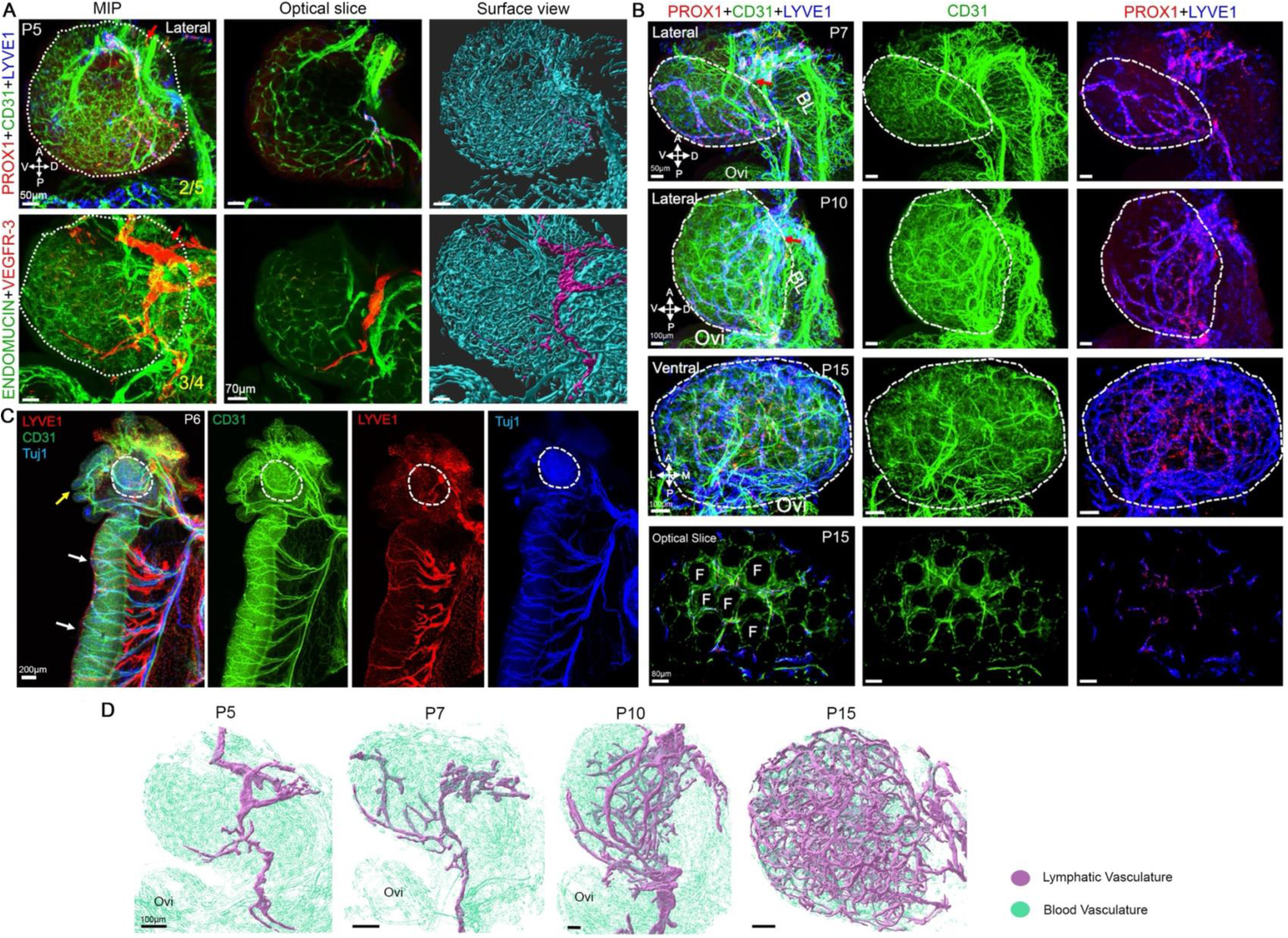
Stepwise postnatal development of the ovarian lymphatic vasculature. High-resolution confocal images of whole-mount ovaries immunostained for various markers at postnatal days P5, P6, P7, P10, and P15. (A) PROX1⁺/LYVE1⁺/VEGFR3⁺ lymphatic vessels first enter the ovary through the hilum (red arrow) at P5 and progressively extend toward the posterior end. White dotted lines outline the ovaries. (B) With advancing development, additional lymphatic vessels sprout from both the hilum and from pre-existing intraovarian vessels. By P15, a lymphatic network is established in close proximity to ovarian follicles. (C) Schematic diagram illustrating the sequential steps of ovarian lymphatic vascular development. (D) Whole-mount immunostaining showing LYVE1⁺ lymphatic vessels traveling in close association with CD31⁺ blood vessels and TUJ1⁺ nerves as they enter the ovary and uterus through the hilum. Red, yellow, and white arrows indicate the ovarian hilum, oviduct, and uterus, respectively. White dotted lines outline the ovaries. Abbreviations: MIP, maximal intensity projection; A, anterior; P, posterior; V, ventral; D, dorsal; BL, broad ligament; Ovi, oviduct. Scale bars and number of animals as indicated. Fractions indicate the number of animals displaying the representative phenotype out of total animals examined.

### Ovarian lymphatic vessels arise from pre-existing lymphatic endothelial cells

Analysis of neonatal ovaries immunostained for PROX1, CD31, and LYVE1 revealed two distinct populations of PROX1⁺ cells that closely associate with CD31⁺ blood capillaries: PROX1⁺/LYVE1⁻ cells (white arrows) and PROX1⁺/LYVE1⁺ cells (yellow arrows; **Figure 2A**). This observation raised the possibility that, similar to lymphangiogenesis in the embryonic dermis (38, 39), ovarian lymphatic vessels might partially arise from capillary-derived endothelial cells that upregulate PROX1, bud from blood vessels, and subsequently acquire LYVE1 expression. To directly determine the origin of ovarian lymphatic endothelial cells (LECs), we performed genetic lineage tracing using TgProx1CreERT2;Rosa26mT/mG mice (40, 41). Tamoxifen was administered at P1 and P3 to label PROX1-expressing cells present prior to ovarian lymphatic invasion, and ovaries were collected at P15 for analysis (**Figure 2B**). Immunostaining for GFP, PROX1, and LYVE1 demonstrated that all PROX1⁺/LYVE1⁺ intraovarian lymphatic vessels were GFP⁺, indicating that they originate from pre-existing PROX1⁺ LECs that are present prior to lymphatic expansion (**Figure 2C**). In addition to GFP⁺/PROX1⁺/LYVE1⁺ lymphatic vessels, two additional GFP-labeled populations were identified: GFP⁺/PROX1⁺/LYVE1⁻ vessels (white arrowheads) and GFP⁺/PROX1⁻/LYVE1⁻ mesenchymal-like cells (yellow arrowheads; **Figure 2C**). The GFP⁺/PROX1⁺/LYVE1⁻ vessels were contiguous with LYVE1⁺ lymphatics through clusters of GFP^hi^/PROX1^hi^ cells, reminiscent of developing lymphatic valve structures. These vessels likely represent lymphatic pre-collectors connected to LYVE1^+^ lymphatic capillaries via lymphatic valves. The GFP⁺/PROX1⁻/LYVE1⁻ population may represent LEC-derived cells that have downregulated canonical lymphatic markers during vascular remodeling; however, whether this population is ovary-specific and functionally significant remains to be determined. Collectively, these findings demonstrate that ovarian lymphatic vessels form through sprouting and expansion of pre-existing LECs, rather than *de novo* differentiation from blood capillaries.

**Figure 2.**
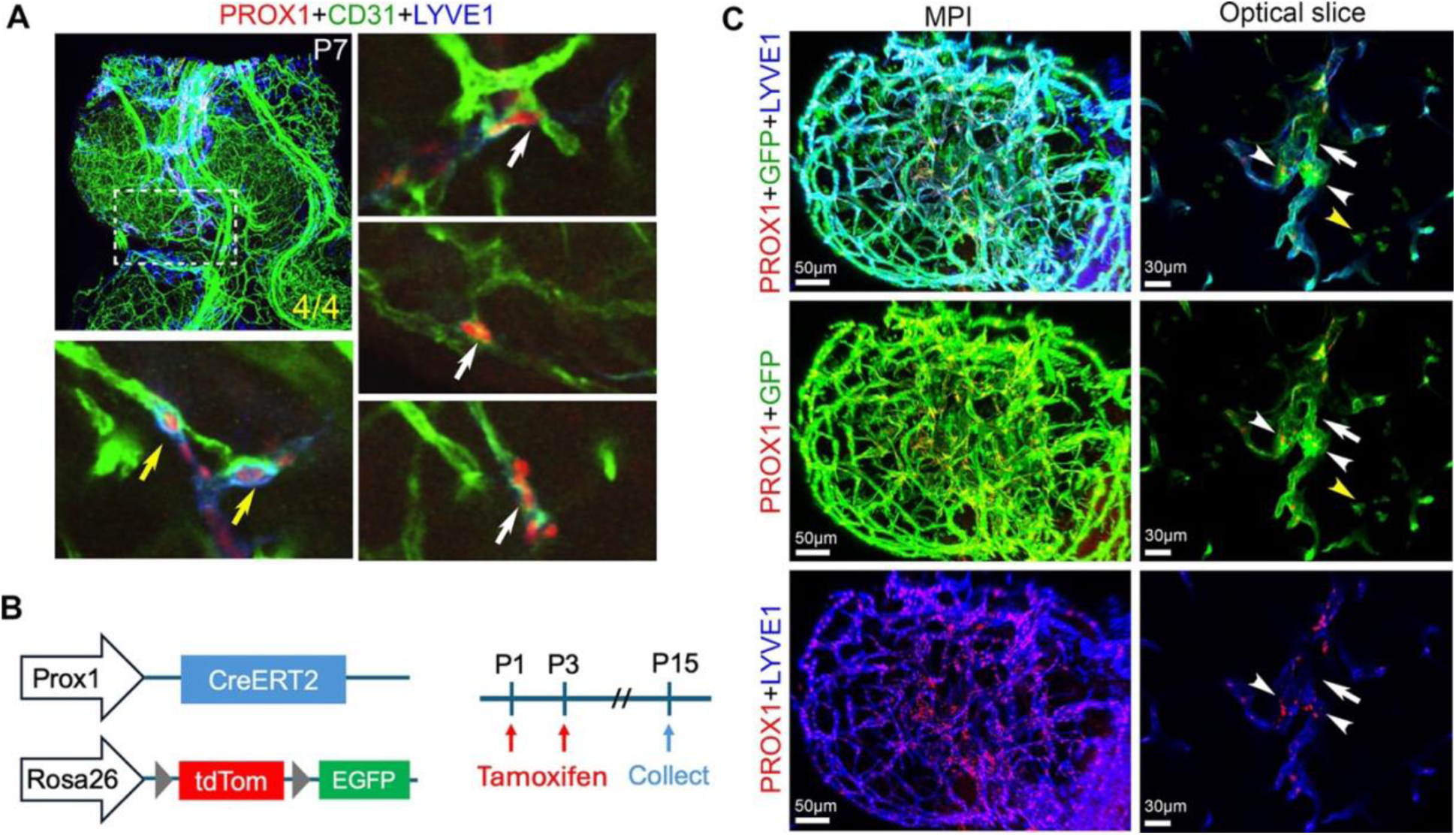
Ovarian lymphatic vasculature originates from pre-existing lymphatic endothelial cells. (A) P7 ovaries immunostained for PROX1, CD31, and LYVE1. White arrows indicate PROX1⁺/LYVE1⁻ cells; yellow arrows indicate PROX1⁺/LYVE1⁺ double-positive cells. Both populations are closely associated with CD31⁺ blood capillaries. (B) Schematic of lineage-tracing experimental design. *TgProx1CreERT2;Rosa26mT/mG* pups received 100 µg tamoxifen at P1 and P3 to label pre-existing PROX1⁺ lymphatic endothelial cells, and ovaries were collected at P15 for analysis. (C) P15 *TgProx1CreERT2;Rosa26mT/mG* ovaries immunostained for GFP, PROX1, and LYVE1. The PROX1⁺/LYVE1⁺ intraovarian lymphatic vasculature is GFP⁺, demonstrating derivation from pre-existing LECs. White arrowheads indicate GFP⁻/PROX1⁻ cell clusters. Yellow arrowheads indicate GFP⁺/PROX1⁻/LYVE1⁻ mesenchymal-like cells. White arrow marks a region of GFP⁺/PROX1⁺/LYVE1⁻ vessel. Scale bars as indicated. N≥4 for each experiment. Fractions indicate the number of animals displaying the representative phenotype out of total animals examined.

### The ovarian lymphatic vasculature consists of hierarchically organized lymphatic capillaries and pre-collecting vessels with functional intraluminal valves

To characterize the structural organization of the mature ovarian lymphatic vasculature, we performed immunofluorescence and ultrastructural analyses of ovaries from 2-month-old Tg(Prox1-GFP) reporter mice, in which GFP is specifically expressed in lymphatic endothelial cells (42). Confocal imaging of GFP, PROX1, and LYVE1 identified two morphologically and molecularly distinct vessel subtypes. GFP⁺/PROX1⁺/LYVE1⁺ lymphatic capillaries were predominately localized to the ovarian periphery and connected at discrete junctions to GFP⁺/PROX1⁺/LYVE1⁻ pre-collecting vessels in the center region (yellow arrowheads; **Figure 3A**). At these junctions, clusters of cells with elevated GFP and PROX1 expression, consistent with lymphatic valves, were observed (white arrowheads; **Figure 3A**,). GFP⁺ lymphatic vessels formed a network surrounding the corpus luteum (CL) but did not penetrate it (**Figure 3B**), confirming the previous observation that the CL constitutes a lymphatic-excluded microenvironment (43, 44). Immunostaining for alpha-smooth muscle actin (SMA), PROX1, LYVE1, CD31, and the valve marker NEOGENIN1 further resolved vessel hierarchy (45). SMA⁺ vascular smooth muscle coverage was detected around both CD31⁺ arteries (yellow arrowhead; **Figure 3C**) and PROX1⁺/CD31⁺ pre-collecting vessels (yellow arrow; **Figure 3C**), whereas LYVE1⁺/PROX1⁺/CD31⁺ capillaries (white arrowheads; **Figure 3C**) connected to LYVE1⁻/PROX1⁺/CD31⁺ collecting-type vessels (white arrows; **Figure 3C**). NEOGENIN1⁺/PROX1⁺ valve structures were identified within PROX1⁺/CD31⁺ pre-collecting vessels (red arrows; **Figure 3C**). To directly assess valve ultrastructure, we performed correlative fluorescence and scanning electron microscopy on 300-µm ovarian sections, as previously described (46) (**Figure 3D**). GFP^hi^ cell clusters identified by fluorescence microscopy (blue and red arrows; **Figure 3D**) corresponded to endothelial cells oriented perpendicular to the vessel axis, forming bileaflet intraluminal valve structures characteristic of functional lymphatic valves. A schematic summary of the ovarian lymphatic vascular architecture is shown in **Figure 3E**.

**Figure 3.**
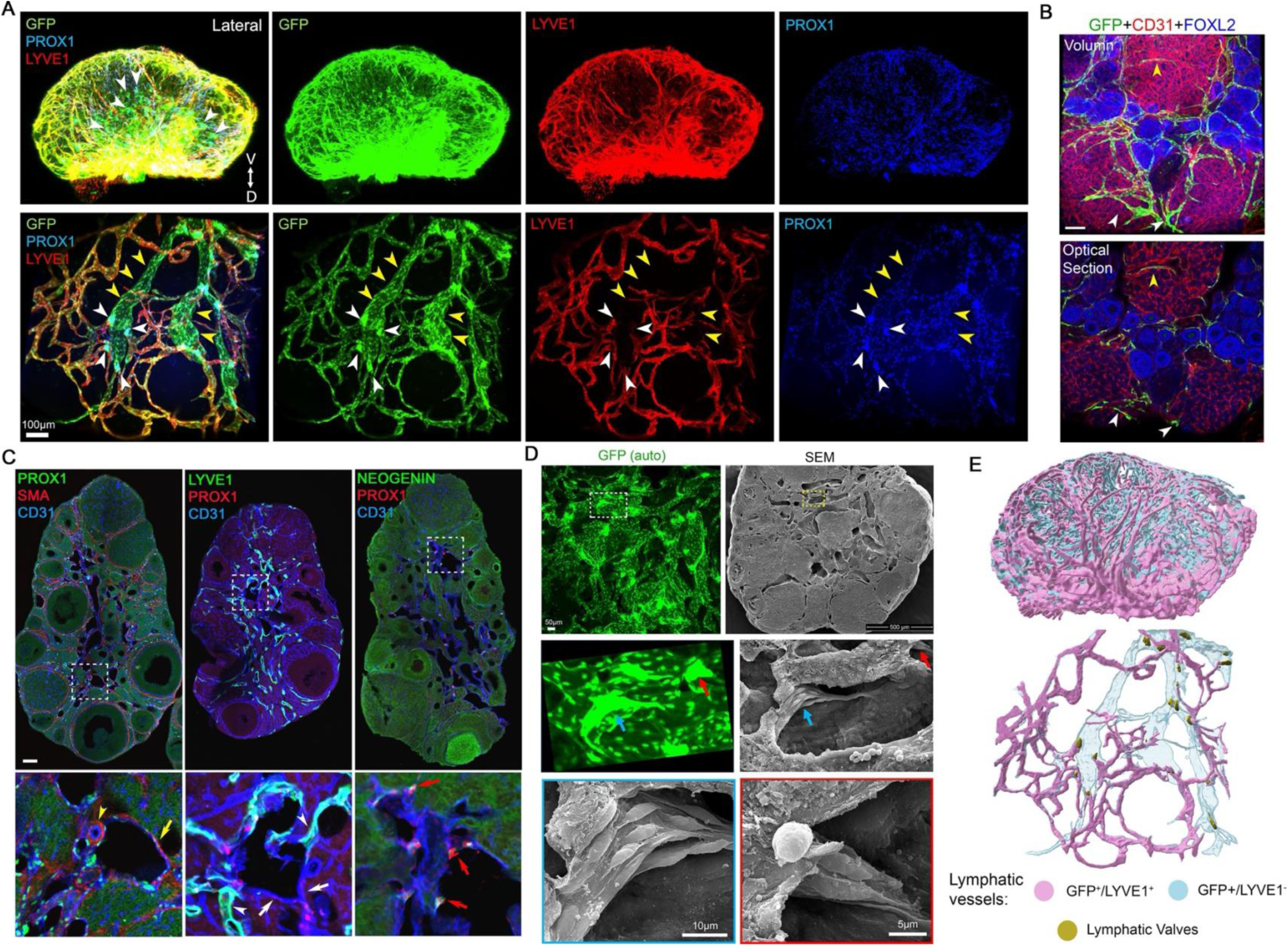
The ovarian lymphatic vasculature consists of lymphatic capillaries and pre-collecting vessels with functional intraluminal valves. (A) Confocal images of 2-month-old *Tg(Prox1-GFP)* ovaries immunostained for GFP, PROX1, and LYVE1. GFP⁺/PROX1⁺/LYVE1⁺ lymphatic capillaries connect to GFP⁺/PROX1⁺/LYVE1⁻ pre-collecting vessels (yellow arrowheads). GFP^hi^/PROX1^hi^ valve-forming cell clusters are present at connection points (white arrowheads). (B) GFP⁺ lymphatic vessels surround the corpus luteum (CL) without penetrating it. (C) Immunostaining of tissue sections from 2-month-old ovaries for SMA, PROX1, LYVE1, CD31, and NEOGENIN1. SMA⁺ smooth muscle cells surround a CD31⁺ artery (yellow arrowhead) and a PROX1⁺/CD31⁺ pre-collecting vessel (yellow arrow). LYVE1⁺/PROX1⁺/CD31⁺ capillaries (white arrowheads) connect to LYVE1⁻/PROX1⁺/CD31⁺ vessels (white arrow). NEOGENIN1⁺/PROX1⁺ lymphatic valves are present in PROX1⁺/CD31⁺ pre-collecting vessels (red arrows). (D) Correlative fluorescence and scanning electron microscopy (SEM) of a 300 µm section of a 2-month-old *Tg(Prox1-GFP)* ovary. GFP^hi^ clusters (blue and red arrows) visible by fluorescence correspond in SEM to cells aligned perpendicular to the direction of flow forming two valve leaflets. (E) Schematic representation of the hierarchical architecture of the ovarian lymphatic vasculature. Scale bars are 50mm or as indicated. N≥4 for each experiment.

### VEGF-C/VEGFR-3 signaling is required for ovarian lymphangiogenesis

Signaling via VEGF-C and VEGFR-3 is the principal pathway governing lymphangiogenesis (47, 48). VEGF-C is required for the sprouting of lymphatic endothelial progenitors from embryonic veins, while postnatal lymphatic expansion is primarily driven by VEGFR-3-mediated proliferation of lymphatic endothelial cells (LECs) derived from pre-existing vessels (49). To determine whether VEGF-C signaling regulates ovarian lymphatic development, we examined the spatial and temporal expression of *Vegfc*, the canonical ligand for the lymphangiogenic receptor VEGFR-3, using RNAscope *in situ* hybridization combined with immunofluorescent staining. *Vegfc* mRNA was detected in stromal cells and CD31⁺ endothelial cells at all postnatal stages examined, including P6, P12, and 2-month-old ovaries (**Figure 4A**). At P6, fluorescence intensity analysis revealed a spatial gradient, with higher *Vegfc* signal at the posterior end of the ovary (blue arrowhead) compared with the anterior end (red arrowhead), consistent with the direction of lymphatic vessel growth. By P12, *Vegfc* transcripts were distributed in stromal and endothelial cells surrounding developing follicles, and a similar perifollicular pattern persisted in 2-month-old ovaries. In addition, low levels of *Vegfc* expression were detected in luteal cells within the corpus luteum (**Figure 4A**). To complement these spatial data with single-cell resolution, we analyzed our published scRNA-seq dataset from 3-month-old mouse ovaries (**Figure 4B–D and Supplementary Figure 1**). *Vegfc* expression was predominantly enriched in stroma A and endothelial cell clusters, with arterial endothelial cells exhibiting the highest transcript levels among all populations (**Figure 4B–D**).

**Figure 4.**
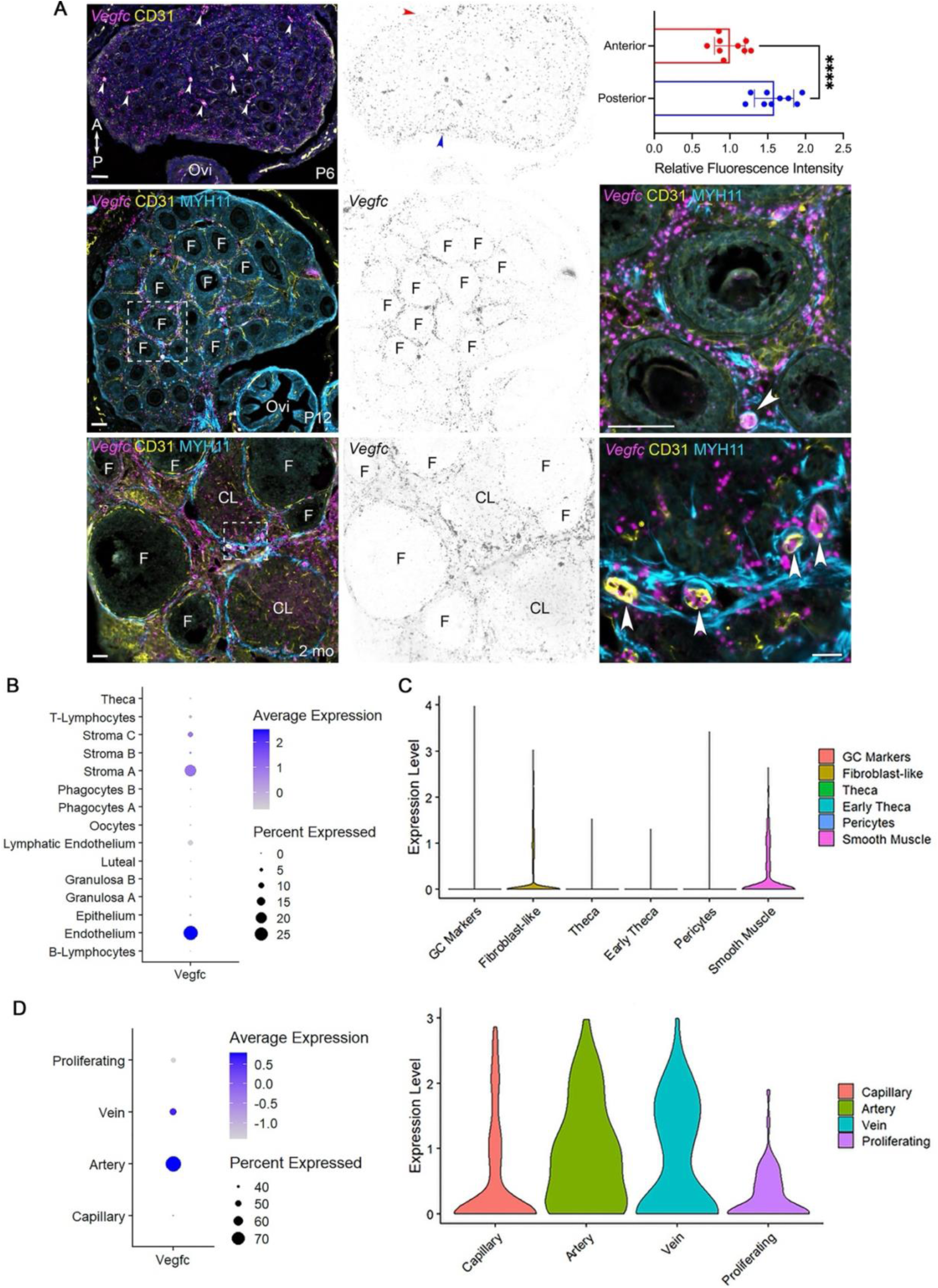
*Vegfc* expression pattern during ovarian lymphatic growth. (A) RNAscope fluorescence in situ hybridization for *Vegfc* on sections of P6, P12, and 2-month-old ovaries. *Vegfc* mRNA is detected in stromal cells and CD31⁺ endothelial cells at all stages. At P6, fluorescence intensity is higher at the posterior end (blue arrowhead) than the anterior end (red arrowhead) of the ovary. At P12, *Vegfc* transcripts are distributed in stromal and endothelial cells surrounding follicles; a similar pattern with additional weak signal in luteal cells is observed at 2 months. (B–D) Single-cell RNA sequencing analysis of 3-month-old mouse ovaries. *Vegfc* is predominantly expressed in stroma A and endothelial cell clusters (B–C), with arterial endothelial cells showing the highest expression levels among all ovarian cell types (D). Statistics: (A) n=3 animals; each dot represents the mean fluorescent intensity within a region of interest (ROI), with 3 sections analyzed per animal. Data are presented as mean ± SD. Statistical significance was determined by two tailed unpaired Student’s *t* test; *\*\*\*\*P < 0.0001*.

Missense mutations in the tyrosine kinase domain of VEGFR-3 account for about approximately 70% of cases of autosomal dominant primary congenital lymphedema known as Milroy disease (47, 50). The *Vegfr3^+/Chy^* mouse, which carries an inactivating point mutation in *Vegfr3*, exhibits chylous ascites and hypoplastic lymphatics and serves as a well-established model of primary lymphedema (51, 52). To directly assess the functional requirement for VEGFR-3 signaling in ovarian lymphangiogenesis, we analyzed ovaries from *Vegfr3^⁺/Chy^* mice, which harbor an inactivating point mutation in the VEGFR-3 kinase domain. At P21-P28, LYVE1⁺ lymphatic vessels were markedly reduced or entirely absent in *Vegfr3^⁺/Chy^* pups, which also exhibited severe chylous ascites. In contrast, CD31⁺ blood vasculature was comparable between wild-type and mutant ovaries (**Figure 4E**). A similarly profound and persistent lymphatic deficiency was observed in the ovaries of surviving *Vegfr3^⁺/Chy^* mice at 4 months of age, with no evidence of compensatory recovery (**Figure 4F**).

To further confirm the requirement for VEGF-C/VEGFR-3 signaling in ovarian lymphatic development, we analyzed *Vegfc^+/-^*mice, which are viable but exhibit lymphatic defects and develop lymphedema due to *Vegfc* haploinsufficiency (53). Consistent with this, *Vegfc^+/−^* ovaries displayed a similar but less penetrant phenotype, with fewer ovaries exhibiting severe lymphatic defects (**Supplementary Figure 2**). Collectively, these findings demonstrate that *Vegfc* is dynamically and spatially regulated during ovarian lymphatic development, forming gradients that guide lymphatic vessel invasion, growth and patterning. Importantly, VEGF-C/VEGFR-3 signaling is both dosage-sensitive and essential for ovarian lymphangiogenesis.

### Lymphatic insufficiency in *Vegfr3^⁺/Chy^* mice elicits hallmarks of ovarian aging

To determine the long-term consequences of ovarian lymphatic insufficiency on reproductive function, we performed a comprehensive analysis of ovarian aging parameters in *Vegfr3^⁺/Chy^* mice and wild-type controls at 7.5 months of age. Staining with Sudan Black B (SBB), a marker of lipofuscin accumulation that reflects cellular oxidative damage, revealed no significant difference in SBB-positive area between control and *Vegfr3^⁺/Chy^* ovaries (**Figure 5A**). This finding indicates that lymphatic insufficiency does not induce oxidative damage in the ovary. In contrast, Picrosirius red staining for fibrillar collagen revealed a significant increase in collagen deposition in *Vegfr3^⁺/Chy^* ovaries compared to wild-type controls (**Figure 5B**), indicating progressive fibrotic remodeling of the ovarian stroma. Ovary-to-body weight ratios were comparable between genotypes (**Figure 5C**), indicating that the phenotype does not reflect gross organ atrophy. Histological quantification of follicle populations revealed a significant reduction in total follicle number, most prominently among tertiary follicles, in 7.5-month-old *Vegfr3^⁺/Chy^*ovaries (**Figure 5D**). Notably, follicle numbers were not significantly different between genotypes at 4 months of age (**Figure 5E**), indicating that the declines in follicular reserve is progressive and age-dependent rather than developmental. Longitudinal estrous cycle monitoring over a 12-day period revealed that *Vegfr3^⁺/Chy^*females spent significantly more time in the estrus phase and less time in the diestrus phase compared with wild-type controls (**Figure 5F–G**), indicating disrupted estrous cyclicity consistent with impaired ovarian endocrine function. Consistent with increased fibrosis and altered cyclicity, *Vegfr3^⁺/Chy^* mice exhibited a delayed onset of pregnancy compared to controls, with wild-type females achieving ∼70–75% pregnancy by day 4, whereas *Vegfr3^⁺/Chy^* mice reaching comparable levels only at later time points (Figure 5H). Collectively, these findings demonstrate that chronic lymphatic insufficiency drives progressive ovarian dysfunction and promotes features commonly associated with ovarian aging, including fibrosis, disrupted cyclicity, and accelerated follicle depletion.

**Figure 5.**
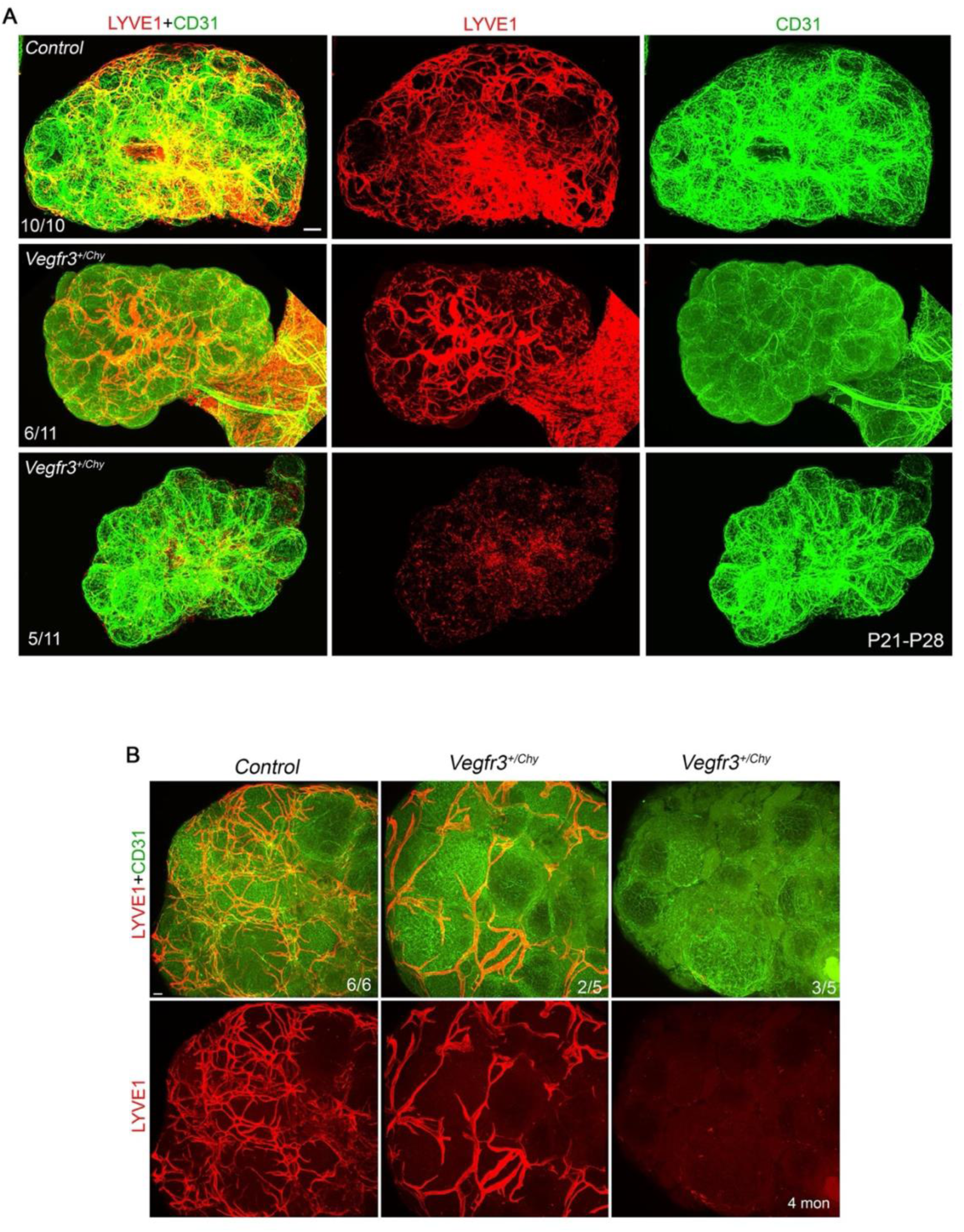
VEGF-C/VEGFR-3 signaling is required for ovarian lymphangiogenesis. (A) Immunostaining for LYVE1 and CD31 in P21–P28 wild-type and *Vegfr3*^⁺/Chy^ ovaries. LYVE1⁺ lymphatic vessels are severely reduced or absent in *Vegfr3*^⁺/Chy^ pups with severe chylous ascites; CD31⁺ blood vasculature is comparable between genotypes. (B) Similar lymphatic deficiency is observed in surviving *Vegfr3*^⁺/Chy^ ovaries at 4 months of age. Scale bars are 50mm. The number of animals analyzed are indicated on the image.

### Lymphatic insufficiency drives immune cell infiltration and myofibroblast activation in the ovary

To investigate the cellular mechanisms underlying lymphatic insufficiency-induced ovarian aging, we performed a detailed immunohistological analysis of immune and stromal cell populations in 7.5-month-old *Vegfr3^⁺/Chy^* and wild-type control ovaries. Staining for cleaved caspase-3 (CC3), FOXL2, and CD31 revealed no significant difference in the percentage of atretic follicles between genotypes (**Figure 6A**), confirming that increased apoptosis of follicles does not account for the observed reduction in follicle numbers. In contrast, the area of SMA⁺ myofibroblasts was significantly increased in *Vegfr3^⁺/Chy^* ovaries, consistent with active fibroblast-to-myofibroblast trans-differentiation, thereby driving collagen deposition (**Figure 6B**). CD3 immunostaining revealed a significant increase in the number of T cells in *Vegfr3^⁺/Chy^*ovaries relative to wild-type controls, indicating enhanced adaptive immune infiltration (**Figure 6C**). Staining for MHCII (a marker of antigen-presenting cells), GPNMB (a marker of multinucleated giant cells, MNGCs), and COLIV (a marker of basement membrane, used to delineate corpus luteum boundaries and blood vessels) revealed that a marked expansion of MHCII⁺ antigen presenting cells (APCs) within ovarian stroma and corpora lutea of *Vegfr3^⁺/Chy^* mice (**Figure 6D**). APCs were frequently observed in aggregates with GPNMB⁺ MNGCs (yellow arrows and arrowheads; **Figure 6D**), and the number of APCs within these aggregates negatively correlated with GPNMB fluorescence intensity, suggesting active phagocytic clearance of GPNMB⁺ cellular debris by APCs (**Figure 6D**). Based on these findings, we propose a mechanistic model in which lymphatic deficiency results in the accumulation of biological waste products in the ovarian interstitium; this waste is processed by resident phagocytes and APCs, which signal to the lymph node and activate T cells; the recruited T cells promote fibroblast-to-myofibroblast differentiation; myofibroblast-derived collagen I increases ovarian stiffness; and elevated stromal stiffness ultimately impairs follicle maturation and reduces follicle number, driving premature ovarian aging (**Figure 6E**).

**Figure 6.**
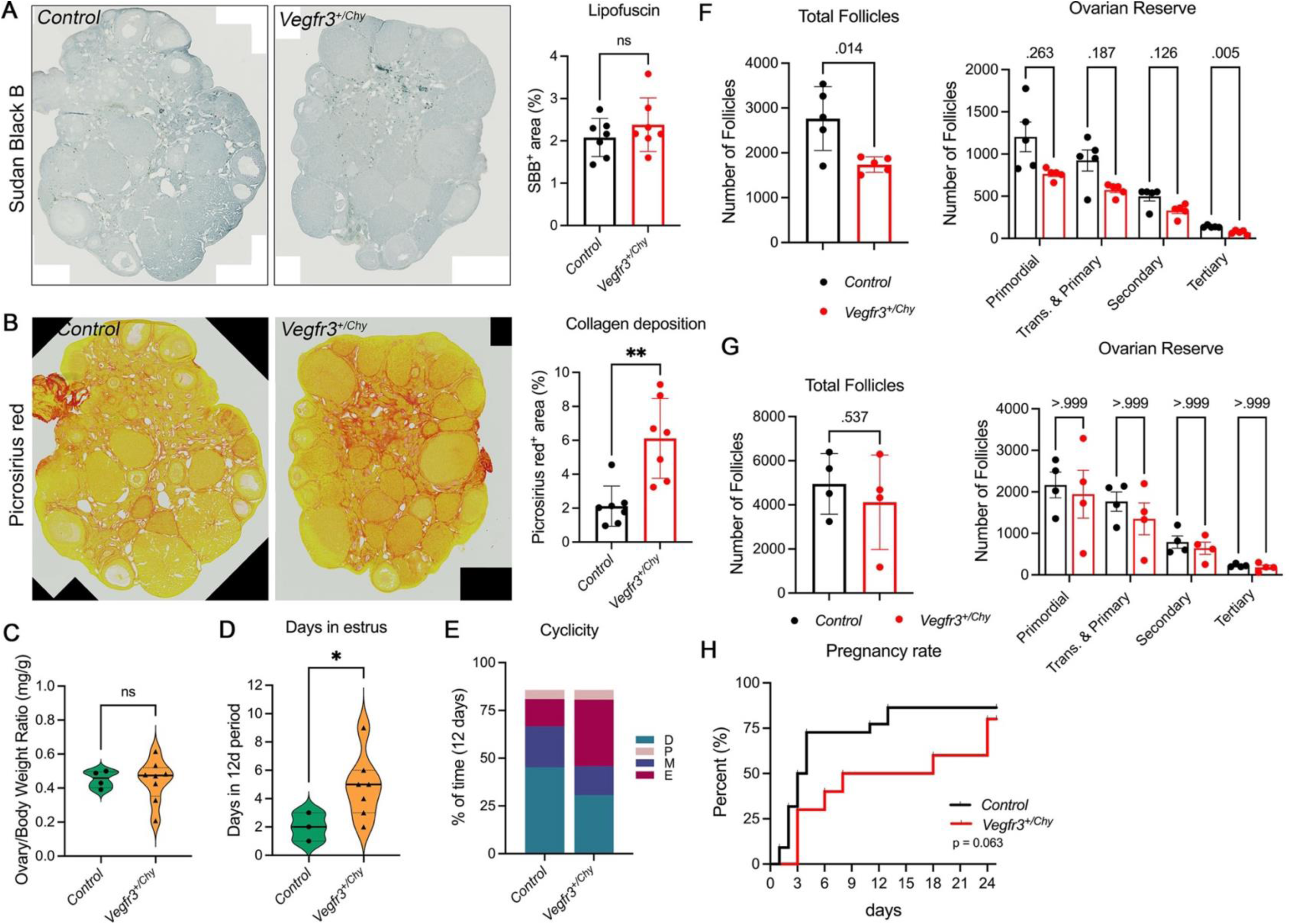
*Vegfr3^⁺/Chy^* ovaries exhibit premature reproductive aging. (A) Sudan Black B (SBB) staining for lipofuscin on sections of 7.5-month-old wild-type and *Vegfr3^⁺/Chy^* ovaries. Quantification shows no significant difference in SBB-positive area between genotypes (right). (B) Picrosirius red staining for fibrillar collagen on sections of 7.5-month-old ovaries. The Picrosirius red–positive area is significantly increased in *Vegfr3^⁺/Chy^* ovaries. Representative images (left) and quantification (right) are shown. (C) Ovary-to-body weight ratio is not significantly different between wild-type and *Vegfr3^⁺/Chy^* mice. (D) Histological quantification of follicle populations in 7.5-month-old ovaries. Total follicle number and tertiary follicle number are significantly reduced in *Vegfr3^⁺/Chy^* ovaries. (E) Follicle counts at 4 months of age reveal no significant difference between genotypes. (F–G) Estrous cycle monitoring over a 12-day period. *Vegfr3^⁺/Chy^*mice spend significantly more time in estrus (F) and less time in diestrus (G) compared to wild-type controls. (H) Kaplan-Meier curve showing the cumulative percentage of pregnant females over a 24-day mating period in control (black) and *Vegfr3^+/Chy^* mice (red). Control animals reached a pregnancy rate of ∼75%, with the majority conceiving within the first 4 days. *Vegfr3^+/Chy^* females showed a delayed onset and reduced cumulative pregnancy rate (∼75% by day 24). Statistics: Data are presented as mean ± SD. Statistical analysis was performed using an unpaired Student’s *t* test; *P* values are indicated on the graphs.

**Figure 7.**
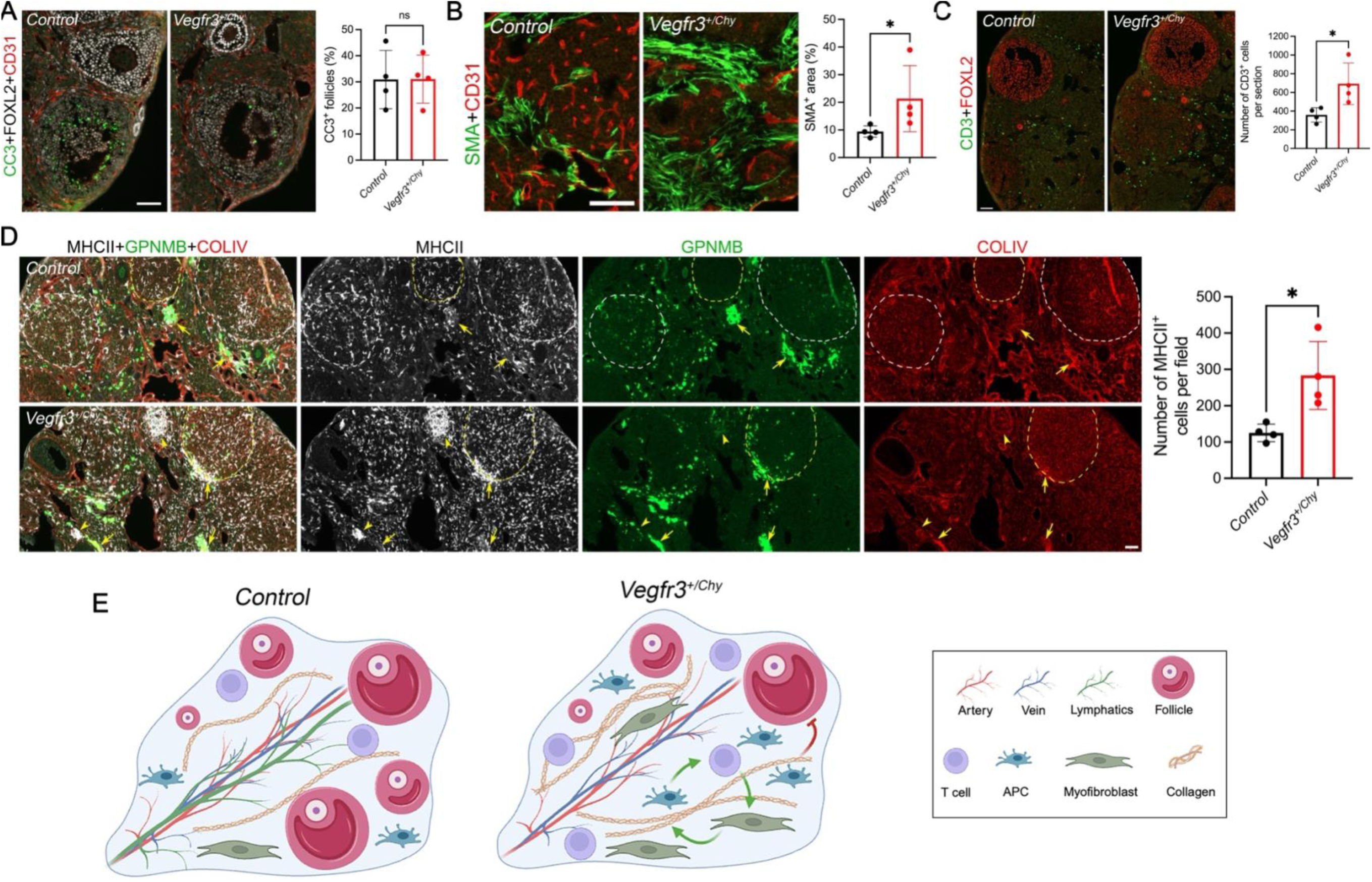
Increased T cells, antigen-presenting cells, and myofibroblasts in *Vegfr3^⁺/Chy^* ovaries. (A) Sections of 7.5-month-old ovaries immunostained for cleaved caspase-3 (CC3), FOXL2, and CD31. The percentage of atretic follicles is not significantly different between wild-type and *Vegfr3^⁺/Chy^* ovaries. (B) SMA immunostaining. The SMA⁺ myofibroblast area is significantly increased in *Vegfr3^⁺/Chy^* ovaries. (C) CD3 immunostaining. The number of CD3⁺ T cells is significantly increased in *Vegfr3^⁺/Chy^* ovaries. (D) Sections of 7.5-month-old ovaries immunostained for MHCII, GPNMB, and COLIV. MHCII⁺ APCs are markedly increased in the stroma and corpora lutea (delineated by dense COLIV⁺ blood vasculature, outlined with white dotted lines) of *Vegfr3^⁺/Chy^*ovaries. APCs aggregate with GPNMB⁺ multinucleate giant cells (yellow arrows and arrowheads). APC number within aggregates negatively correlates with GPNMB fluorescence intensity, indicating active phagocytic clearance of GPNMB⁺ material. (E) Proposed mechanistic model: lymphatic deficiency leads to accumulation of biological waste in the ovarian interstitium; waste activates APCs, which signal to lymph nodes and recruit T cells; T cells promote myofibroblast differentiation; myofibroblast-secreted collagen I increases ovarian stiffness; elevated stiffness impairs follicle maturation and reduces follicle number, driving premature ovarian aging. Data are presented as mean ± SD. *p < 0.05; ns, not significant. Statistical analysis was performed using an unpaired Student’s *t* test.

## DISCUSSION

In this study, we present the first systematic characterization of the postnatal development and physiological function of the lymphatic vasculature in the mouse ovary. Using high-resolution 3D imaging, genetic lineage tracing, tissue-clearing, *in situ* RNA detection, and single-cell transcriptomics, we define the precise timing and route of lymphatic invasion into the ovary, establish the cellular origin of ovarian lymphatics, and identify VEGF-C/VEGFR-3 signaling as the essential driver of ovarian lymphangiogenesis. Critically, we demonstrate that chronic lymphatic insufficiency in *Vegfr3^⁺/Chy^* mice promotes the emergence of hallmarks commonly observed in the aging ovary, including reduced follicular reserve, increased fibrosis, and elevated immune cell infiltration. Together, these findings establish the ovarian lymphatic vasculature as an indispensable component of the reproductive microenvironment and define a previously unrecognized pathway through which lymphatic insufficiency contributes to ovarian dysfunction.

The temporal relationship between ovarian follicle development and the postnatal appearance of lymphatic vessels in the ovary is a finding of considerable developmental significance. Our data show that lymphatic vessels first invade the ovary at P5 through the hilum and progressively extend inward, reaching the mid-ovarian region by P10 and forming a network in proximity to developing follicles by P15. This timeline closely parallels the progression of folliculogenesis: primordial follicles are established during late fetal and early neonatal life, completing formation around postnatal day 4, with the first wave of follicle activation and primary follicle formation occurring during the first two postnatal weeks in mice (54, 55). The near-simultaneous arrival of lymphatics and the expansion of the growing follicle pool suggests that the lymphatic vasculature may be recruited, at least in part, in response to the increasing metabolic and immunological demands imposed by the follicular microenvironment. Conversely, follicular growth factors or the stromal remodeling that accompanies folliculogenesis may provide the pro-lymphangiogenic cues that guide lymphatic invasion. It remains unknown if this temporal coordination reflects a causal relationship by which early perturbations of follicle activation alter the pace or extent of lymphatic development. The finding that lymphatic deficiency does not detectably impair follicle numbers at 4 months of age, but does so by 7.5 months of age, suggests that the initial follicular reserve is established independently of lymphatic support, whereas sustained folliculogenesis and follicle maintenance across reproductive life depend on an intact lymphatic network.

Our RNAscope and scRNA-seq data establish that *Vegfc* is expressed throughout ovarian postnatal development in stromal cells and vascular endothelium, with arterial endothelial cells showing the highest expression at all stages examined. This pattern is consistent with the known requirement for a paracrine VEGF-C gradient to induce sprouting, as VEGF-C is essential for the budding of endothelial cells committed to the lymphatic lineage, a gradient whose disruption blocks lymphatic formation entirely (47). The graded *Vegfc* expression along the anterior-to-posterior axis of the P6 ovary with higher levels at the posterior end suggests that a spatial VEGF-C gradient may direct lymphatic growth from the hilum toward the posterior pole. The near-complete absence of LYVE1⁺ lymphatics in *Vegfr3^⁺/Chy^*ovaries at both P21–P28 and 4 months of age, without a corresponding defect in CD31⁺ blood vasculature, is consistent with the previous reports of normal blood vasculature in Chy mice and the established postnatal specificity of VEGFR-3 signaling for lymphatic rather than blood endothelium (51, 56).

At 7.5 months, *Vegfr3^⁺/Chy^* mice exhibit features of accelerated ovarian aging, including increased collagen deposition, reduced tertiary and total follicle numbers, an abnormal estrous cyclicity with prolonged estrus. In addition, *Vegfr3^⁺/Chy^* females display a delayed onset of pregnancy compared to controls, which may reflect impaired ovulatory timing likely due to stromal fibrosis and altered cyclicity. Together, these findings indicate that lymphatic insufficiency accelerates ovarian aging rather than inducing acute follicular toxicity. Consistently, ovarian fibrosis driven by extracellular matrix accumulation and age-associated increases in collagen with reduced hyaluronan is a hallmark of reproductive aging (16, 18). The unchanged atresia rate and cleaved caspase-3⁺ granulosa cell fraction further argue against a direct pro-apoptotic effect, instead implicating stromal remodeling as a key driver of impaired follicle maturation. Notably, the stromal fibrosis observed in *Vegfr3^⁺/Chy^* ovaries resembles the pathology of primary lymphedema, where impaired lymphatic function induces chronic inflammation, immune cell infiltration, fibroblast activation, and progressive collagen deposition (57–59). The involvement of shared cellular mediators including SMA⁺ myofibroblasts, MHCII⁺ antigen-presenting cells, and CD3⁺ T cells suggests that ovarian fibrosis represents a tissue-specific manifestation of a broader lymphedema-associated fibrotic program. These observations highlight potential translational opportunities, as anti-inflammatory and anti-fibrotic strategies effective in lymphedema models may be applicable to ovarian lymphatic insufficiency and raises the possibility that subclinical lymphatic dysfunction could predispose to accelerated ovarian fibrosis and premature follicular decline (58, 60–62).

These findings carry several important implications. First, they establish a novel functional role for organ-specific lymphatics: the ovarian lymphatic vasculature is not simply a passive fluid-draining conduit, but an active regulator of the immune and stromal microenvironment required for sustained folliculogenesis and reproductive longevity. Second, they raise the clinically relevant question of whether subtle lymphatic dysfunction contributes to idiopathic premature ovarian insufficiency (POI) in women, a condition in which ovarian follicles are depleted and cease to function normally both as reproductive and endocrine organs in women under 40 years old, and which is characterized by deficient ovarian sex hormones and decreased follicles that accelerate the onset of menopause (63). Lymphatic abnormalities have not been systematically evaluated in POI patients, and our data provide a rationale for doing so. Third, the identification of the VEGF-C/VEGFR-3 axis as essential for ovarian lymphatic maintenance raises the possibility that VEGF-C-based pro-lymphangiogenic therapies, currently in clinical and preclinical development for lymphedema (64, 65), might eventually be explored as a strategy to preserve ovarian function in at-risk populations. Several limitations merit acknowledgment. The *Chy* mutation causes systemic lymphatic insufficiency, and while the ovarian phenotype we describe is internally consistent and mechanistically coherent, formal ovary-specific lymphatic ablation experiments will be required to definitively rule out contributions from systemic lymphedema or altered hormonal axes. Additionally, while our data define a clear VEGF-C/VEGFR3 requirement for ovarian lymphangiogenesis, the upstream regulators that establish the spatiotemporal *Vegfc* gradient during ovarian development and whether FSH, estrogen, or other reproductive hormones dynamically regulate ovarian lymphangiogenesis across the estrous cycle remain important open questions for future investigation.

## METHODS

### Mice

Tg(Prox1-CreERT2), Rosa26mT/mG, Tg(Prox1-GFP), *Vegfc^+/CreERT2^* and *Vegfr3^+/Chy^* mouse lines were described previously (40–42, 66). The Tg(Prox1-CreERT2) and Tg(Prox1-GFP) lines were obtained through material transfer agreement (MTA) with the investigators who originally developed these models. C57BL/6J, HET3 and Rosa26mT/mG mice were purchased from Jackson Laboratory. *Vegfr3^+/Chy^* line was obtained from European Mouse Mutant Archive (EMMA). *Vegfr3^+/Chy^*and *Vegfc^+/CreERT2^* were maintained on a mixed C57BL/6J;NMRI;HET3 background. All other lines were maintained on C57BL/6J:NMRI mixed background. All mice were housed on a standard chow diet. For lineage tracing experiment, pups were fed with 5ml of tamoxifen (20mg/ml) on P1 and P3.

### Antibodies

A complete list of antibodies is provided in Supplementary Methods.

### Immunofluorescent (IF) Staining

Whole mount IF staining was performed as previously described using iDISCO protocol (67). Neonatal ovaries were collected together with the oviduct and a portion of the uterine horns following euthanasia by asphyxiation, and fixed overnight in 2% paraformaldehyde (PFA) at 4 °C. After PBS washes, the bursa and periovarian adipose tissue were removed. Adult animals were perfused with cold PBS followed by 4% PFA. Reproductive tissues (ovary, oviduct and partial uterine horns) were collected, post fixed overnight in 2% PFA, and the ovaries were subsequently dissected from surrounding tissues. After staining, samples were cleared in Ce3D clearing solution at room temperature until optically transparent(68). Cleared tissues were mounted between two coverslips with spacers and imaged using a Nikon AX R confocal microscope. Image analysis was performed using Imaris software. IF on sections was performed as previously described (69).

### Correlative Fluorescence and Scanning Electron Microscopy

Correlative fluorescence and SEM was performed according to our previous protocol (46, 69).

### RNAscope Combined with Immunofluorescence

RNAscope *in situ* hybridization was performed on 12 μm cryosections using the RNAscope 2.5 High Definition (HD) Detection Kit (RED) (Advanced Cell Diagnostics, Cat# 322360), according to the manufacturer’s instructions with minor modifications. Briefly, slides were baked at 60°C for 1 hour, fixed in 4% PFA for 10 minutes, and treated with Proteinase IV for 30 minutes, followed by hydrogen peroxide treatment for 10 minutes. *Vegfc* probes were hybridized for 2 hours at 40°C, and signals were amplified through sequential amplification steps (AMP1–4) provided in the kit. Sections were then incubated with Fast RED substrate, followed by immunofluorescence staining after washing, as previously described. Images were acquired using a Nikon C2 confocal microscope and analyzed with ImageJ software.

### scRNAseq Analyses

The scRNA-seq dataset utilized in this study was originally generated and described by Isola et al. (2024) (70–72). We adhered to the Data processing and cell-type annotations established in the primary publication. Top-level cell classifications and previously defined sub-clusters for stroma and theca populations were directly adopted from the published ovarian atlas. To further resolve the lymphatic compartment, the endothelial cell population was subset and subjected to high-resolution reclustering, followed by annotation based on the expression of canonical marker genes. Data visualizations, including dot plots and violin plots were generated using the Seurat R package. All secondary data processing followed the quality control and normalization parameters established in the original publication.

### Histological Analyses

Histological analyses were conducted on ovaries from 4-7 animals per group per time-point. One ovary from each animal was excised and fixed in 4% paraformaldehyde containing 1.5% glutaraldehyde, processed, and serially sectioned. To evaluate ovarian reserve, one in six serial sections covering the whole ovary was stained with hematoxylin and eosin (H&E) and follicles at all developmental stages (primordial, primary, secondary, and antral) were counted as previously described (70–72). Picrosirius Red staining was used to evaluate collagen deposition, and Sudan Black B staining to measure lipofuscin accumulation. For both assays, a randomly selected mid-ovarian section was analyzed from each animal and quantified as reported previously (70–72).

### Estrus Cycle Assessments

Estrus cycle stages were determined by vaginal cytology using a non-invasive lavage method described previously (73). In brief, mice were gently restrained, and 20 µL of sterile 1× PBS was flushed into the vaginal canal. The recovered fluid was placed on a glass slide, smeared, and air-dried. The slides were stained using a modified Romanowsky staining kit (Hema 3™, Fisher Scientific) following manufacturer’s instructions. Vaginal smears were collected daily for 12 consecutive days. Estrous cycle stages (proestrus, estrus, metestrus, and diestrus) were determined based on the relative proportions of nucleated epithelial cells, cornified epithelial cells, and leukocytes observed under light microscopy.

### Statistics

All data are presented as mean ± SD. For comparisons between two independent groups with n ≥ 5 per group, statistical significance was determined using a two-tailed unpaired Student’s *t* test. For comparisons between two groups with n ≤ 4 per group, the Mann-Whitney U test was applied. The cumulative pregnancy rate was performed using the Kaplan-Meier method, and differences between curves were assessed using the Gehan-Breslow-Wilcoxon test. A *P* value of less than 0.05 was considered statistically significant. All statistical analyses and graphical representations of quantitative data were performed using GraphPad Prism software (version 10.0; GraphPad Software).

### Study Approval

All mice were housed and handled according to the institutional IACUC protocols of Oklahoma Medical Research Foundation.

### Data Availability

The datasets generated through this work are available upon reasonable request from the corresponding author.

## AUTHOR CONTRIBUTIONS

X.G. and M.B.S. conceived the project and designed the experiments. S.B. and X.G. performed the majority of the experiments with contributions from L.C., J.S., and R.S.S. S.B. and X.G. analyzed data and wrote the manuscript. M.B.S. L.X. and R.S.S. provided resources and edited the manuscript. All authors approved the final version of the manuscript.

## ACKNOWLEDGMENTS

The authors thank Taija Makinen for providing Tg(Prox1-CreERT2) mice and Young Kwon Hong for providing Tg(Prox1-GFP) mice. We also thank Amber English for assistance with Imaris imaging analysis, Lisa Whitworth and Brent Johhson for assistance with SEM imaging, Harini Bagavant for helpful discussions and Zoltan Ungvari for insightful comments on the manuscript. During the preparation of this manuscript, Claude and ChatGPT (GPT4) were used for editing assistance, specifically to improve language and readability.

