## supplementary materials for "Development and Function of Ovarian Lymphatic Vasculature"

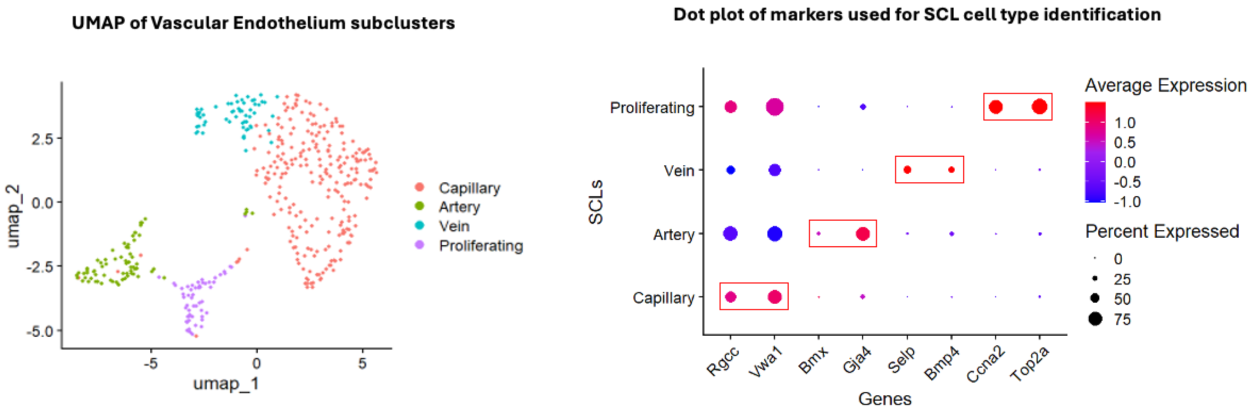


**Supplementary Figure 1. Single-cell transcriptomic analysis of vascular endothelial subclusters.** (A) UMAP embedding of vascular endothelial cells reveals four transcriptionally distinct subclusters: capillary (salmon), artery (green), vein (teal), and proliferating endothelial cells (lavender). (B) Dot plot displaying the average expression and percent of cells expressing canonical marker genes used to assign cell type identity to each subclustered cell lineage (SCL). Dot size reflects the percentage of cells within each SCL expressing the indicated gene; dot color indicates average normalized expression level (red, high; blue, low). Red boxes highlight the marker genes most discriminatory for each SCL identity. Capillary endothelial cells are marked by *Rgcc* and *Vwa1*; arterial endothelial cells by *Bmx* and *Gja4*; venous endothelial cells by *Selp*; and proliferating endothelial cells by *Ccna2* and *Top2a*.

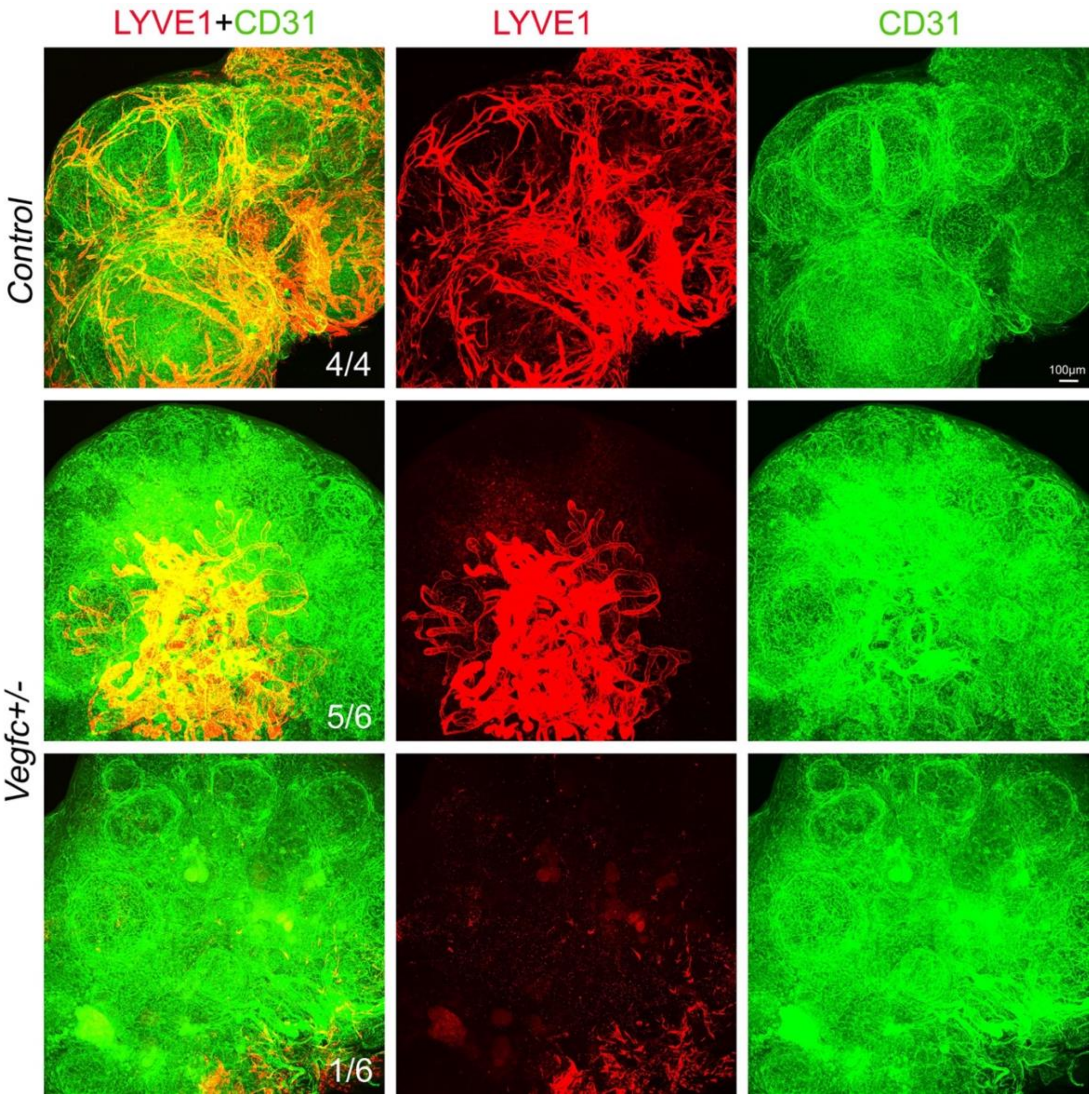


**Supplemental Figure 2. Lymphatic vessel development in control and *Vegfc* heterozygous embryos.** Whole-mount immunofluorescence staining of embryonic tissues showing co-labeling with the lymphatic endothelial marker LYVE1 (red) and the pan-endothelial marker CD31 (green). Representative images are shown for control (top row, 4/4), *Vegfc*+/− embryos with a mild phenotype (middle row, 5/6), and *Vegfc*+/− embryos with a severe phenotype (bottom row, 1/6). Controls display an elaborately branched lymphatic network with extensive LYVE1-positive vessels throughout the ovary. In contrast, *Vegfc*+/− embryos show a progressive reduction in lymphatic vessel complexity and coverage, with the most severely affected ovary exhibiting markedly diminished LYVE1 signal and failure of lymphatic network elaboration despite preserved CD31-positive blood vasculature. Fractions indicate the number of animals displaying the representative phenotype out of total animals examined. Scale bar as indicated.

| **Antibody** | **Manufacture** | **Catalog Number** |
|---|---|---|
| GFP | abcam | ab13970 |
| VEGFR3 | R&D SYSTEMS | AF743 |
| CD31 | BD Pharmingen | 553370 |
| LYVE1 | R&D SYSTEMS | AF2125 |
| CD3 | Biolegend | 100202 |
| COLIV | abcam | ab19808 |
| GPNMB | R&D SYSTEMS | AF2330 |
| MHCII | Invitrogen | 14-5321-82 |
| CC3 | R&D SYSTEMS | AF835 |
| MYH11 | Novus Biologicals | NBP2-66967 |
| NEOGENIN1 | Novus Biologicals | AF1097 |
| FOXL2 | Novus Biologicals | NB100-1277 |
| Tuj1 | abcam | ab18207 |
| Cy3-conjugated Donkey Anti-Goat IgG (H+L) | Jackson ImmnunoReserach Lab | 705-165-147 |
| Cy3-conjugated Donkey Anti-Rabbit IgG (H+L) | Jackson ImmnunoReserach Lab | 711-165-152 |
| Cy5-conjugated Donkey Anti-Rat IgG (H+L) | Jackson ImmnunoReserach Lab | 712-175-150 |
| Cy5-conjugated Donkey Anti-Goat IgG (H+L) | Jackson ImmnunoReserach Lab | 705-175-147 |
| Cy5-conjugated Donkey Anti-Rabbit IgG (H+L) | Jackson ImmnunoReserach Lab | 711-175-152 |
| Alexa Fluor 488-conjugated Donkey Anti-Rabbit IgG (H+L) | Jackson ImmnunoReserach Lab | 711-547-003 |
| Alexa Fluor 488-conjugated Donkey Anti-Goat IgG (H+L) | Jackson ImmnunoReserach Lab | 705-545-147 |
| Alexa Fluor 488-conjugated Donkey Anti-Chicken IgG (H+L) | Jackson ImmnunoReserach Lab | 703-545-155 |
| Alexa Fluor 488-conjugated Donkey Anti-Rat IgG (H+L) | Jackson ImmnunoReserach Lab | 712-545-153 |
| Cy3-conjugated Anti $\alpha$-Smooth Muscle Actin (SMA) | Sigma | C6198 |
| FITC-conjugated Anti $\alpha$-Smooth Muscle Actin (SMA) | Sigma | F3777 |
